# DolphinNext: A graphical user interface for creating, deploying and executing Nextflow pipelines

**DOI:** 10.1101/689539

**Authors:** Onur Yukselen, Osman Turkyilmaz, Ahmet Rasit Ozturk, Manuel Garber, Alper Kucukural

## Abstract

The emergence of high throughput technologies that produce vast amounts of genomic data, such as next-generation sequencing (NGS) are transforming biological research. The dramatic increase in the volume of data makes analysis the main bottleneck for scientific discovery. The processing of high throughput datasets typically involves many different computational programs, each of which performs a specific step in a pipeline. Given the wide range of applications and organizational infrastructures, there is a great need for a highly-parallel, flexible, portable, and reproducible data processing frameworks. Flexibility ensures that pipelines can support a variety of applications without requiring one-off modifications. Portability ensures that users can leverage computationally available resources and work within economic constraints. Reproducibility warrants credibility to the results and is particularly challenging in the face of the sheer volume of data and the complexity of processing pipelines that vary widely between users.

Several platforms currently exist for the design and execution of complex pipelines (e.g. Galaxy, GenePattern, GeneProf). Unfortunately, these platforms lack the necessary combination of parallelism, portability, flexibility and/or reproducibility that are required by the current research environment. To address these shortcomings, Nextflow was implemented to simplify portable, scalable, and reproducible scientific pipelines using containers. We have used Nextflow capabilities as leverage and developed a user interface, DolphinNext, for creating, deploying, and executing complex Nextflow pipelines for high throughput data processing. The guiding principle of DolphinNext is to facilitate the building and deployment of complex pipelines using a modular approach implemented in a graphical interface. DolphinNext provides: 1. A drag and drop user interface that abstracts pipelines and allows users to create pipelines without familiarity in underlying programming languages. 2. A user interface to monitor pipeline execution that allows the re-initiation of pipelines at intermediate steps 3. Reproducible pipelines with version tracking and stand-alone versions that can be run independently. 4. Seamless portability to distributed computational environments such as high-performance clusters or cloud computing environments.

## INTRODUCTION

Analysis of high-throughput data is now widely regarded as the major bottleneck in modern biology(1). In response, resource allocation has dramatically skewed towards computational power, with significant impacts on budgetary decisions(2). One of the complexities of high-throughput sequencing data analysis is that a large number of different steps are often implemented with a heterogeneous set of programs with vastly different user interfaces. As a result, even the simplest sequencing analysis requires the integration of different programs and familiarity with scripting languages. Programming was identified early on as a critical impediment to genomics workflows. Indeed, microarray analysis became widely accessible only with the availability of several public and commercial platforms, such as GenePattern(3) and Affymetrix(4), that provided a user interface to simplify the application of a diverse set of methods to process and analyze raw microarray data.

A similar approach to sequencing analysis was later implemented by Galaxy(5), GenomicScape(6), and other platforms(3, 7–14). Each of these platforms has a similar paradigm: Users upload data to a central server and apply a diverse, heterogeneous set of programs through a standardized user interface. As with microarray data, these platforms allow users without any programming experience to perform sophisticated analyses on sequencing data obtained from different protocols such as RNA Sequencing (RNA-Seq) and Chromatin Immunoprecipitation followed by Sequencing (ChIP-Seq) and carry out sophisticated analysis. Users are able to align sequencing reads to the genome, assess differential expression, and perform gene ontology analysis through a unified point and click user interface. A comparison between widely used platforms is shown in Table 1.

**Table 1.** Comparison to related applications.

|  | DolphinNext | Galaxy | Sequanix | Taverna | Arvados |
| --- | --- | --- | --- | --- | --- |
| Platform<sup>a</sup> | JS/PHP | Python | Python | Java | Go |
| Workflow management system | Nextflow | Galaxy | Snakemake | Taverna | Arvados |
| Native task support<sup>b</sup> | Yes (any) | No | Yes (bash only) | Yes (bash only) | Yes |
| Common workflow language<sup>c</sup> | No | Yes | No | No | Yes |
| Streaming processing<sup>d</sup> | Yes | No | No | No | Yes |
| Code sharing integration<sup>e</sup> | Yes | No | No | No | Yes (github) |
| Workflow modules<sup>f</sup> | Yes | Yes | Yes | Yes | Yes |
| Workflow versioning<sup>g</sup> | Yes | Yes | No | No | No |
| Automatic error failover<sup>h</sup> | Yes | Yes | No | No | Yes |
| Nested workflows | Yes | Yes | No | Yes | No |
| Used syntax/ semantics | own/own | XML/own | Python/own | own/own | Python/own |
| Web-based | Yes | Yes | No | No | No |
| Container Support |  |  |  |  |  |
| Docker support | Yes | Yes | Yes | No | Yes |
| Singularity support | Yes | Yes | Yes | No | No |
| Built-in batch schedulers |  |  |  |  |  |
| LSF | Yes (Native) | Yes (DRMAA) | Yes (Native) | No | No |
| SGE | Yes (Native) | Yes (DRMAA) | Yes (Native) | Yes (Native) | No |
| SLURM | Yes (Native) | Yes (DRMAA) | Yes (Native) | No | Yes (Native) |
| IGNITE | Yes (Native) | No | No | No | No |
| Built-in cloud |  |  |  |  |  |
| AWS (Amazon Web Services) | Yes | Yes | No | Yes | Yes |
| Autoscaling | Yes | Yes | No | No | Yes |

- The technology and the programming language in which each framework is implemented. - The ability of the framework to support the execution of native commands and scripts without re-implementation of the original processes. - Support for the CWL specification. - Ability to process tasks inputs/outputs as a stream of data. - Support for code management and sharing platforms, such as GitHub. - Support for module, sub-workflows or workflow compositions. - Ability to track pipeline changes and to execute different versions at any point in time. - Support for automatic error handling and resume execution mechanism.

While current platforms are a powerful way to integrate existing programs into pipelines that carry end-to-end data processing, they are limited in their flexibility. Installing new programs is usually only done by administrators or advanced users. This limits the ability of less skilled users to test new programs or simply add additional steps into existing pipelines. This development flexibility is becoming ever more necessary as genome-wide assays are becoming more prevalent and data analysis pipelines becoming increasingly creative (15).

Similarly, computing environments have also grown increasingly complex. Institutions rely on a diverse set of computing options ranging from large servers, higher performance computing clusters, to cloud computing. Data processing platforms need to be easily portable to be used in different environments that best suit the computational needs and budgetary constraints of the project. Further, with the increased complexity of analyses, it is important to ensure reproducible analyses by making analysis pipelines easily portable and less dependent on the computing environment where they were developed (16). Lastly, it is necessary to have a flexible and scalable pipeline platform that can be used both by individuals with smaller sample sizes as well as by medium and large laboratories that need to analyze hundreds of samples a month, or centralized informatics cores that analyze data produced by multiple laboratories.

NextFlow is a recently developed workflow engine built to address many of these needs (17). The NextFlow engine can be configured to use a variety of executors (e.g. SGE, SLURM, LSF, Ignite) in a variety of computing environments. A pipeline that leverages the specific multi-core architecture of a server can be written on a workstation and easily re-used on a high-performance cluster environment (e.g. Amazon and Google cloud) whenever the need for higher parallelization arises. Further, NextFlow allows in-line process definition that simplifies the incorporation of small processes that implement new functionality. Not surprisingly, NextFlow has quickly gained popularity, as reflected by several efforts to provide curated and revisioned NextFlow-based pipelines such as nf-core, Pipeliner (18) and CHIPER (19), which are available from a public repository (20).

In spite of its simplicity, NextFlow can get unwieldy when pipelines become complex, and maintaining them becomes taxing. Here we present DolphinNext, a user friendly, highly scalable, and portable platform that is specifically designed to address current challenges in large data processing and analysis. DolphinNext builds on NextFlow shown in Figure 1. To simplify pipeline design and maintenance, DolphinNext provides a graphical user interface to create pipelines. The pipelines can be exported as NextFlow files or readily run within DolphinNext. Graphical design of workflows is critical when dealing with large and complex workflows. Both advance NextFlow users as well as users with no prior experience benefit from the ability to visualize dependencies, branch points and parallel processing opportunities. DolphinNext goes beyond providing a NextFlow graphical design environment and addresses many of the needs of high-throughput data processing: First, DolphinNext helps with reproducibility enabling the easy distribution and running of pipelines. In fact, reproducible data analysis requires making both the code and the parameters used in the analysis accessible to researchers. To easily enable third parties to reproduce analyses, complete pipelines along with the computing environment used must be available and need to be easily shared(21–24). DolphinNext allows users to package pipelines into portable containers(25, 26) that can be run as stand-alone applications because they include the exact versions of all software dependencies that were tested and used. The automatic inclusion of all software dependencies vastly simplifies the effort needed to share, run and reproduce the exact results obtained by a pipeline.

**Figure 1.**
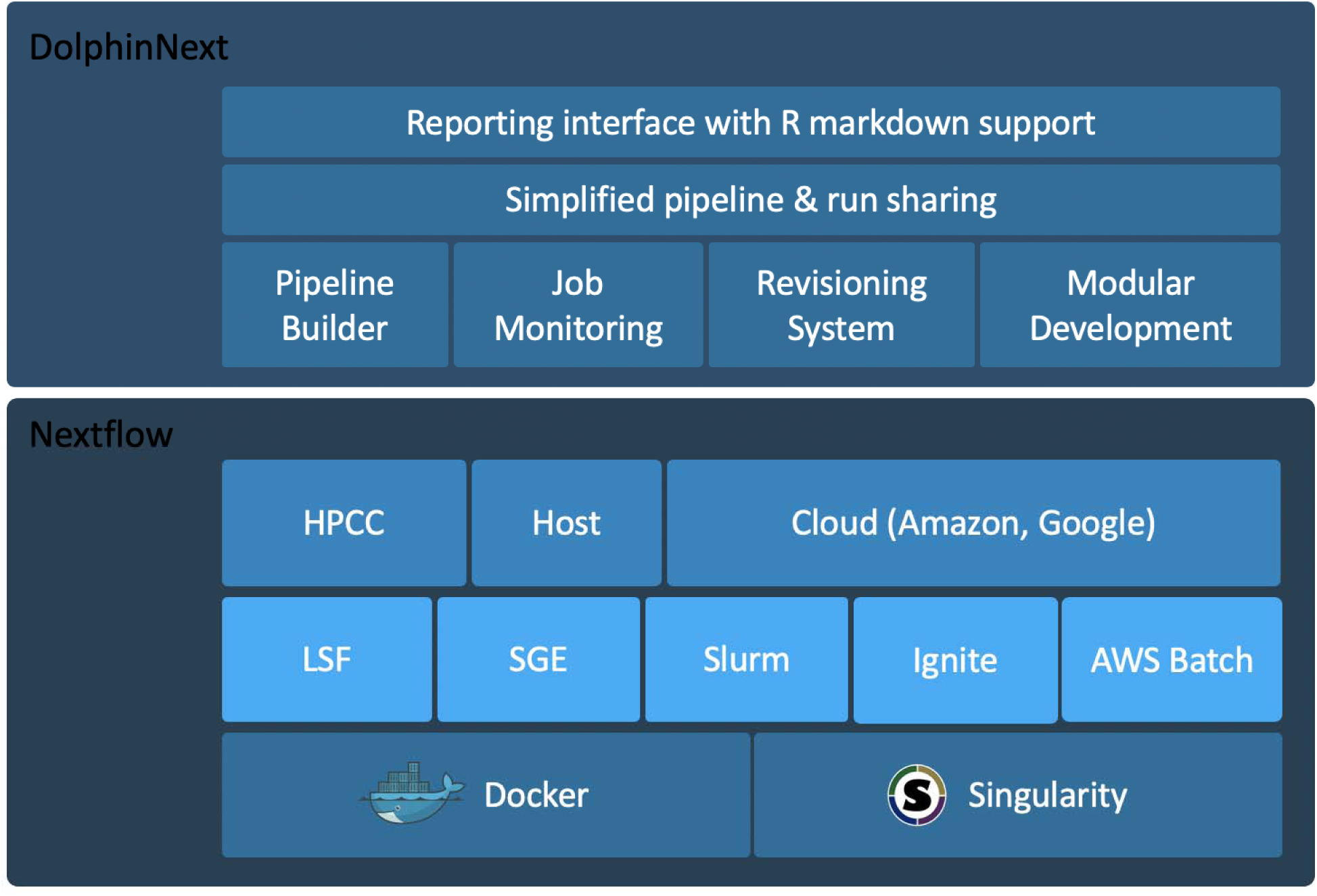
DolphinNext builds on Nextflow and simplifies creating complex workflows.

Second, DolphinNext goes beyond existing data processing frameworks: Rather than requiring data to be uploaded to an external server for processing, DolphinNext is easily run across multiple architectures, either locally or in the cloud. As such it is designed to process data where the data resides rather than requiring users to upload data into the application. Further, DolphinNext is designed to work on large datasets, without needing customization. It can thus support the needs of large sequencing centers and projects that generate vast sequencing data such as ENCODE(27), GTex(28), and TCGA (The Cancer Genome Atlas) Research Network (https://www.cancer.gov/tcga) that have had the need to develop custom applications to support their needs. DolphinNext can also readily support smaller laboratories that generate large sequencing datasets.

Third, as with NextFlow, DolphinNext is implemented as a generic workflow design and execution environment. However, in this report we showcase its power by implementing sequencing analysis pipelines that incorporate best practices derived from our experience in genomics research. This focus is driven by our current use of DolphinNext, but its architecture is designed to support any workflow that can be supported by NextFlow.

Sequence analysis requires an iterative approach that highlights the flexibility of DolphinNext, it consists of two main tasks: data processing and data analysis (29) (**Figure 2**). Data processing involves taking raw sequencing data to produce a count matrix in which features (genes, peaks or any genomic annotation) are scored across samples. Data analysis uses this count matrix to perform comparisons across conditions (differential analysis), unbiased analysis (clustering, principal component analysis), or supervised analysis (data modeling, classifier building)(30). An informative count matrix requires systematic data processing steps that are consistent across samples and even projects. As opposed to data analysis, which involves more ad-hoc, and data driven decisions, data processing can be standardized into a workflow or pipeline that, once established, needs very few changes. In general, data processing requires great computing power and storage needs relative to data analysis. Data processing also greatly benefits from parallelization, and for large projects it is generally carried out on a high-performance computing clusters or cloud computing environments. A typical data processing pipeline can be broken down into three steps: pre-processing, processing, and post-processing. **Pre-processing**: includes eliminating low quality reads or low-quality bases through filtering, read trimming and adapter removal. These steps are critical to improve alignment and assembly operations, which make up the central **processing** step. In general, processing of sequencing reads involves alignment to a genome or reference annotations (e.g. a transcriptome) or an assembly process. **Post-processing** involves evaluating the quality of mapping or assembly before creating a count table, quality checks of alignment and/or assembly steps, and outputs the quantification of genomic features as consolidated count tables. To enable post-processing and data quality assessment, DolphinNext automatically creates genome browser loadable files such as Tile Data Files (TDF) for the Integrative Genome Viewer(31) and Wiggle Track Format (WIG, or BigWIG) for the UCSC genome browser(32, 33). In addition, DolphinNext produces alignment reports that summarize read coverage of genomic annotations (e.g. exonic, intergenic, UTR, intronic) in order to help assess whether the library captured expected genomic regions.

**Figure 2.**
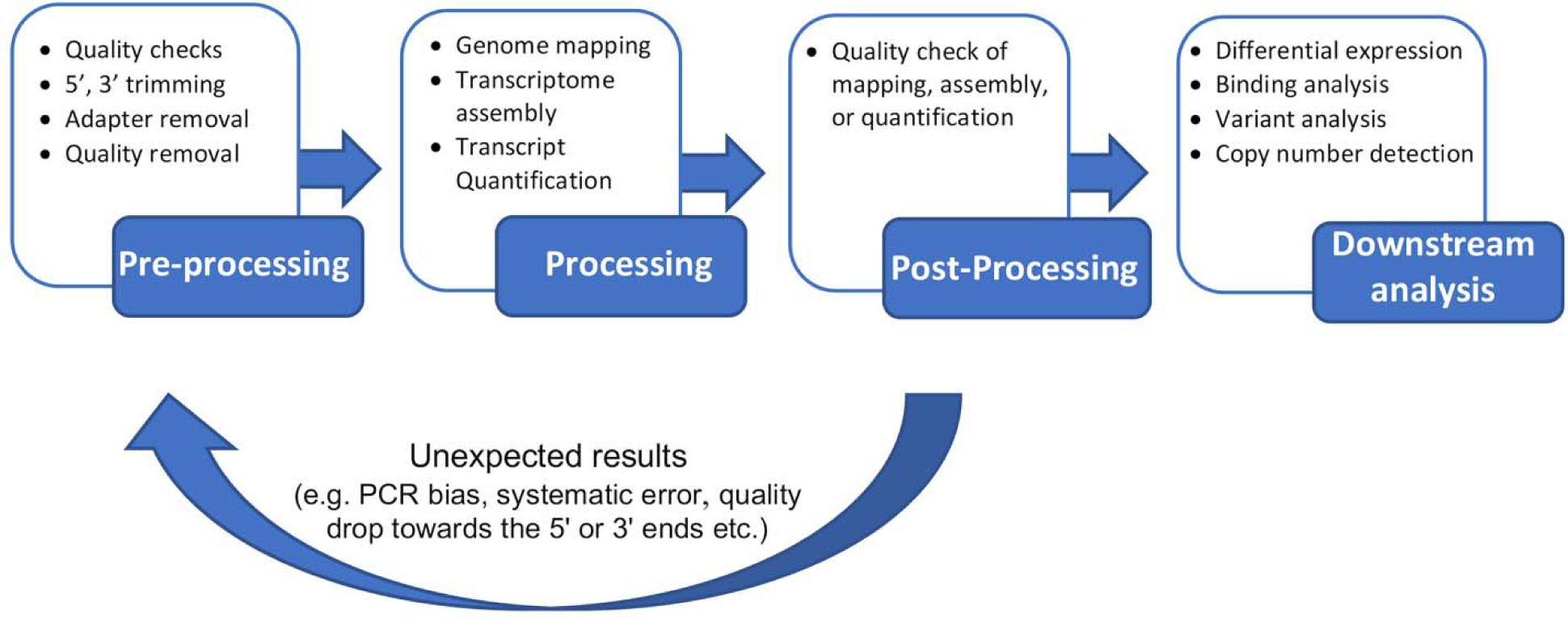
DolphinNext allows for implementation of a complete sequencing analysis cycle.

In conclusion, DolphinNext provides an intuitive interface for weaving together processes each of which have dependent inputs and outputs into complex workflows. DolphinNext also allows users to easily reuse existing components or even full workflows as components in new workflows; in this way, it enhances portability and helps to create more reproducible and easily customizable workflows. Users can monitor job status and, upon identifying errors, correct parameters or data files and restart pipelines at the point of failure. These features save time and decrease costs, especially when processing large data sets that require the use of cloud-based services.

In summary, the key features of DolphinNext include:

- Simple pipeline design interface
- Powerful job monitoring interface
- User specific queueing by job submissions tied to user accounts
- Easy re-execution of pipelines for new sets of samples by copying previous runs
- Simplified sharing of pipelines using the GitHub repository hosting system (github.com)
- Portability across computational environments such as workstations, computing clusters, or cloud-based servers
- Built-in pipeline revision control
- Full access to application run logs
- Parallel execution of non-dependent processes
- Integrated data analysis and reporting interface with R markdown support

Finally, DolphinNext can be easily packaged in portable containers that include the exact programs and versions of the pipelines, making it easy to distribute pipelines (including relevant datasets) to ensure easy replication of computational results.

## MATERIALS AND METHODS

The DolphinNext workflow system, with its intuitive web interface, was designed for a wide variety of users, from bench biologists to expert bioinformaticians. DolphinNext is meant to aid in the analysis and management of a large datasets on High Performance Computing (HPC) environments (e.g LSF, SGE, Slurm Apache Ignite), cloud services, or personal workstations.

DolphinNext is implemented with php, mysql and javascript technologies. At its core, it provides a drag-and-drop user interface for creating and modifying Nextflow pipelines. Nextflow(17) is a language to create scalable and reproducible scientific workflows. In creating DolphinNext, we aimed to simplify Nextflow pipeline building by shifting the focus from software engineering to bioinformatics processes using a graphical interface that requires no programming experience. DolphinNext supports a wide variety of scripting languages (e.g. bash, perl, python, groovy) to create processes. Processes can be used in multiple pipelines, which increases the reusability of the process and simplify code sharing. To that end, DolphinNext supports user and group level permissions so that processes can be shared among a small set of users or all users in the system. Users can repurpose existing processes used in any other pipelines, which eliminates the need to create the same process multiple times. These design features allow users to focus on only their unique needs rather than be concerned with implementation details.

DolphinNext pipelines are versioned, allowing users to improve pipelines or processes, while allowing the previous, stable versions to still be available. All the pipelines and processes in DolphinNext carry unique revision IDs (UIDs) that enable users to run an older version of a pipeline or process should a specific reason to reproduce an old result emerge.

To facilitate reproducibility of data processing and the execution of pipelines in any computing environment, DolphinNext leverages Nextflow’s support for Singularity and Docker container technologies (25, 26). This allows the execution of a pipeline created by DolphinNext to require only Nextflow and a container software (Singularity or Docker) to be installed in the host machine. Containerization simplifies complex library, software and module installation, packaging, distribution and execution of the pipelines by including all dependencies. When distributed with a container, DolphinNext pipelines can be readily executed in remote machines or clusters without the need to manually install third party software programs. Alternatively, DolphinNext pipelines can be exported as Nextflow code and distributed in publications. Exported pipelines can be executed from the command line upon ensuring that all dependencies are available in the executing host. Moreover, multiple executors, clusters, or remote machines can easily be defined in DolphinNext in order to perform computations in any available Linux-based cluster or workstation.

User errors can cause premature failure of pipelines, while also consuming large amounts of resources. Additionally, users may want to explore the impact of different parameters on the resulting data. To facilitate re-running of a pipeline, DolphinNext builds on NextFlow’s ability to record a pipeline execution state, enabling the ability to re-execute or resume a pipeline from any of its steps, even after correcting parameters or correcting a process. Pipelines can also be used as templates to process new datasets by modifying only the dataset specific parameters.

In general, pipelines often require many different parameters, including the parameters for each individual program in the pipeline, system parameters (e.g. paths, commands), memory requirements, and the number of processors to run each step. To reduce the tedious set-up of complex pipelines, DolphinNext makes use of extensive pre-filling options to provide sensible defaults. For example, physical paths of genomes, their index files, or any third-party software programs can be defined for each environment by the admin. When a pipeline uses these paths, the form loads pre-filled with these variables, making it unnecessary to fill them manually. The users still can change selected parameters as needed, but the pre-filling of default parameters speeds up the initialization of a new pipeline. For example, in an RNA-Seq pipeline, if RefSeq annotations(34) are defined as a default option, the user can change it to Ensembl annotations(35) both of which may be located at predefined locations, alternatively the user may specify a custom annotation by supplying a path to the desired annotation file.

Finally, when local computing resources are not sufficient, DolphinNext can also be integrated into cloud-based environments. DolphinNext readily integrates with Amazon AWS where, a new, dedicated computer cluster can easily be set up within DolphinNext with Nextflow’s Amazon cloud support. On AWS, necessary input files can be entered from a shared file storage EFS, EBS, or s3, and output files can also be written on s3 or other mounted drives(36–38).

### General Implementation and Structure

DolphinNext has four modules: The **profile module** specifically designed to support a multi-user environment, allows an administrator to define the specifics of their institutional computing environment, a **pipeline builder** to create reusable processes and pipelines, the **pipeline executor** to run pipelines, and **reports** to monitor the results.

#### 1. Profile Module

Users may have access to a wide range of different computing environments: A workstation, Cloud Computing, or a high-performance computing clusters where jobs are submitted through a job scheduler such as IBM’s LSF, SLURM or Apache Ignite. DolphinNext relies on NextFlow(17) to encapsulate computing environment settings and allows administrators to rely on a single configuration file that enables users to run on diverse environments with minimal impact on user experience. Further, cloud computing and higher performance computing systems keep track of individual user usage to allocate resources and determine job scheduling priorities. DolphinNext supports individual user profiles and can transparently handle user authentication. As a result, DolphinNext can rely on the underlying computing environment to enforce proper resource allocation and fair sharing policies. By encapsulating the underlying computing platform and user authentication, administrators can provide access to different computing architectures, and users with limited computing knowledge can transparently access a vast range of different computing environments through a single interface.

#### 2. Pipeline Builder

While Nextflow provides a powerful platform to build pipelines, it requires advanced programming skills to define pipelines as it requires users to use a programming language to specify processes, dependencies, and the execution order. Even for advanced users, when pipelines become complex, pipeline maintenance can be a daunting task.

DolphinNext facilitates pipeline building, maintenance, and execution by providing a graphical user interface to create and modify NextFlow pipelines. Users choose from a menu of available processes **(Figure 3. A)** from the menu and use drag and drop functionality to create new pipelines by connecting processes through their input and output parameters **(Figure 3. B)**. Two processes can only be connected when the output data type of one is compatible with the input data type of the second **(Figure 3. C)**. Upon connecting two compatible processes DolphinNext creates all necessary execution dependencies. Users can readily create new processes using the process design module (see below). Processes created in the design module are immediately available to the pipeline designer without any installation in DolphinNext.

**Figure 3.**
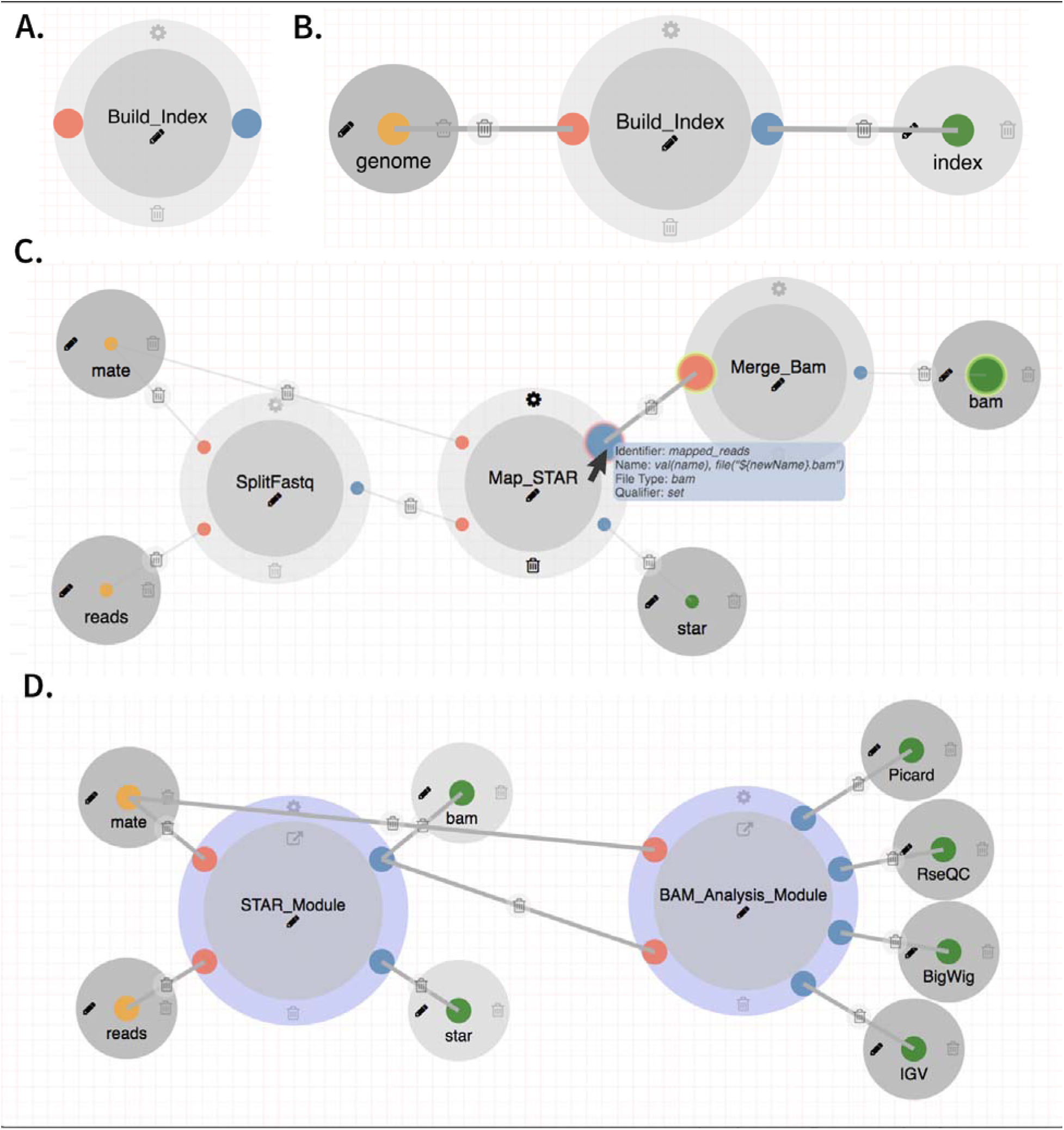
**A.** A process for building index files **B.** Input and output parameters attached to a process **C.** The STAR alignment module connected through input/output with matching parameter types. **D.** The RNA-Seq pipeline can be designed using two nested pipelines: the STAR pipeline and the BAM analysis pipeline.

The UI supports auto-saving to avoid loss of work if users forget to save their work. Once a pipeline is created, users can track revisions, edit, delete and share either as a stand-alone container NextFlow program, or in PDF format for documentation purposes.

The components of the pipeline builder are the process definition module, the pipeline designer user interface, and the revisioning system:

### Process Design Module

Processes are the core units in a pipeline, they perform self-contained and well-defined operations. DolphinNext users designing a pipeline can define processes using a wide variety of scripting languages (e.g. Shell scripting, Groovy, Perl, Python, Ruby, R). Once a process is defined, it is available to any pipeline designer. A pipeline is built from individual processes by connecting outputs with inputs. Whenever two processes are connected, a dependency is implicitly defined whereby a process that consumes the output of another only runs once this output is generated. Since each process may require specific parameters, DolphinNext provides several features to simplify maintenance of processes and input forms that allow the user to select parameters to run them.

#### Automated input form generation

Running all processes within a pipeline requires users to specify many different parameters ranging from specifying the input (e.g. input reads in fastq format, path to reference files) to process specific parameters (e.g. alignment maximum mismatches, minimum base quality to keep). To gather this information, users fill out a form or set of forms to provide the pipeline with all necessary information to run. The large number of parameters makes designing and maintaining the user interfaces that gather this information time consuming and error prone. DolphinNext includes a meta language that converts defined input parameters to web controls. These input parameters are declared in the header of a process script with the help of additional directives. For example, if a pipeline designer needs to create a dropdown selection method for the genome reference input parameter, she would specify this parameter by adding the directives below:

~~~
params.genome_build = "" //* **@dropdown @options**:"human_hg19, mouse_mm10, custom"
~~~

Similarly, to specify whether the transcript quantification program, RSEM(39) should be run in a pipeline, the pipeline designer can capture this information by specifying a run_RSEM parameter with a default option of running and two possible options ‘yes’ and ‘no’:

~~~
params.run_RSEM = "yes" //* **@dropdown @options**:"yes","no”
~~~

In both cases, these directives ensure that the process form includes the options and defaults intended by the designer, irrespective of the pipeline where the process is included.

#### Form autofill support

The vast majority of users work with default parameters and only need to specify a small fraction of all the parameters used by the pipeline. To simplify pipeline usage, we designed an autofill option to provide sensible process defaults and compute environment information. This allows process builders to include directives and obtain default parameters for general usage, such as RSEM parameters to control the number of threads or to determine whether alignment files will be generated upon completion of the run. In this way, this pipeline can be run with the default parameters. For example, the default parameters for an RSEM process can be set to run with four processors and no alignment files will be generated to reduce the space usage by specifying:

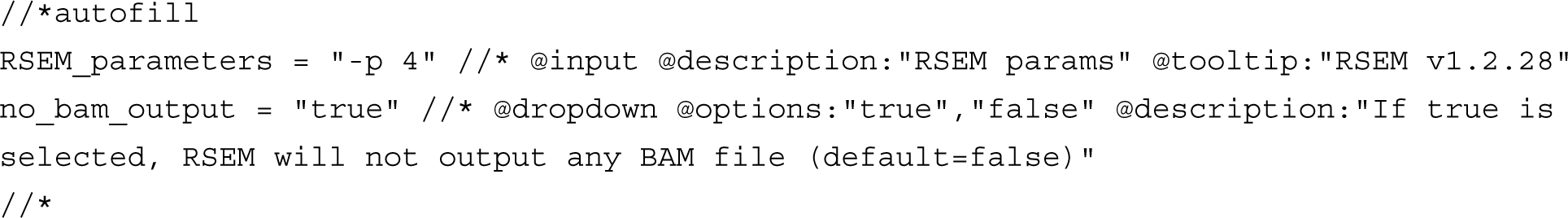

Autofill is meant to provide sensible defaults; however, users can override them as needed. The descriptions of parameters and tooltips are also supported in these directives. Figure 4 shows the description of a defined parameter in RSEM settings.

**Figure 4.**
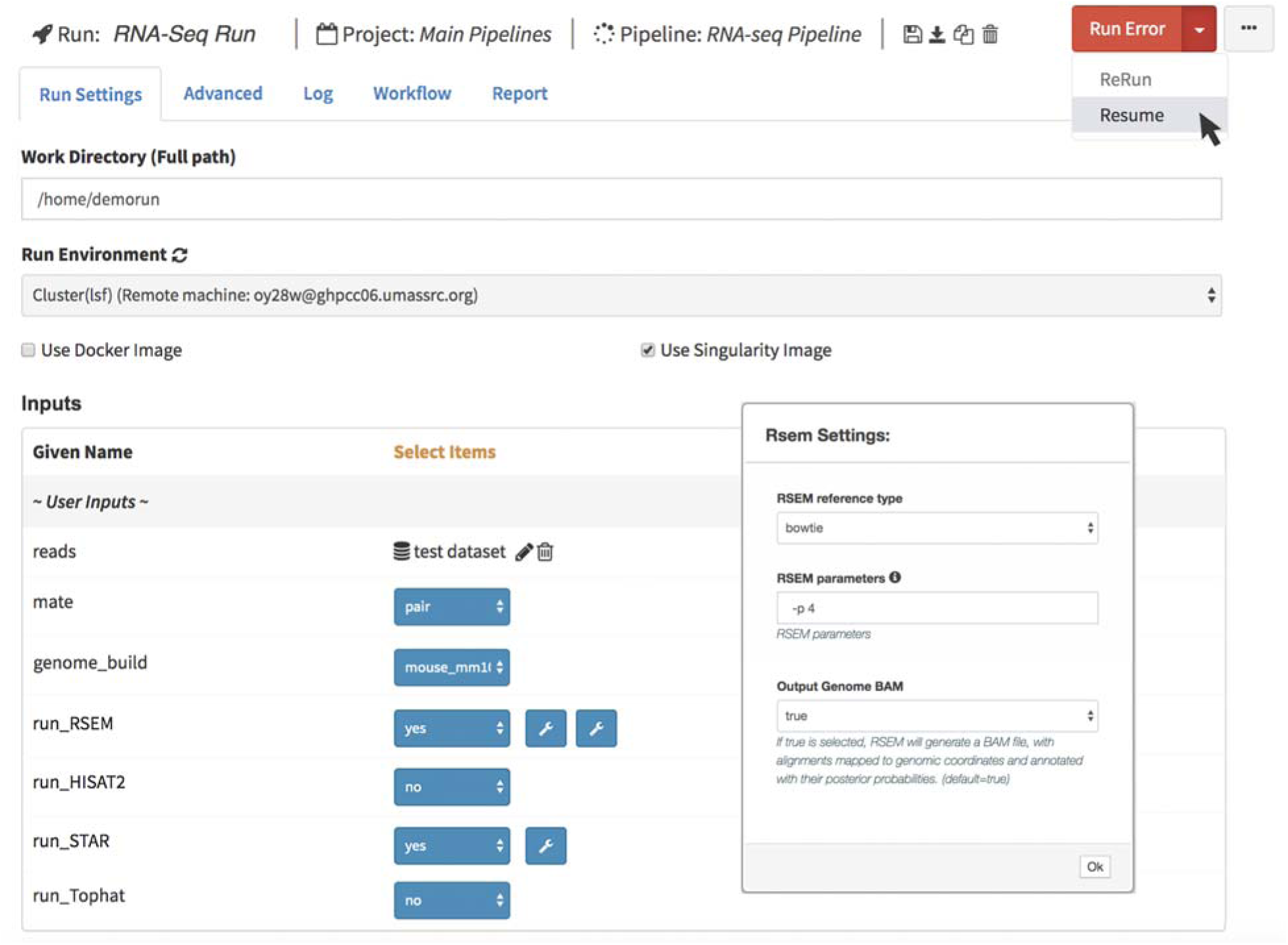
Resuming RNA-Seq pipeline after changing RSEM parameters

### Revisioning, Reusability and Permissions system

DolphinNext implements a revisioning system to track all changes in a pipeline or a process. In this way, all versions of a process or pipeline are accessible and can be used to reproduce old results or to evaluate the impact of new changes. In addition, DolphinNext provides safeguards to prevent the loss of previous pipeline versions. If a pipeline is shared (publicly or within a group), it is not possible to make changes on its current revision. Instead, users must create a new version to make changes. Hence, we keep pipelines safe from modifications yet allowing for improvements to be available in new revisions. DolphinNext uses a local database to assign and store a unique identifier (UID) to every pipeline created and every revision made. A central server may be configured to assigned UIDs across different DolphinNext installations so that pipelines can be identified from the UID, regardless of where they were created. Pipeline designers and users can select any version of a pipeline for execution or editing. In addition to database support, DolphinNext integrates with a GitHub repository so that pipelines can be more broadly shared. DolphinNext can seamlessly push pipelines to a specified repository or branch. In addition to storing the pipeline code, DolphinNext updates its own pipeline or revision database record with the GitHub commit id to keep the revisions that have been synced with a GitHub repository. To support tests and continuous integration of pipelines, we have integrated travis-ci (travis-ci.org), the standard for automated testing. Pipeline designers can define the Travis-ci test description document within the DolphinNext pipeline builder. When a pipeline is updated and pushed to GitHub, it automatically triggers the travis-ci tests. To enable travis-ci automation, pipeline designers specify a container (25, 26) within the pipeline builder.

#### User permissions and pipeline reusability

To increase reusability, DolphinNext supports pipeline sharing. DolphinNext relies on a permissions system similar to that used by the UNIX operating systems. There are three levels of permissions: user, group and world. By default, a new pipeline is owned and only visible to the user who created it. The user can change this default by creating a group of users and designating pipelines as visible to users within that group. Alternatively, the user can make a pipeline available to all users. DolphinNext further supports a refereed workflow by which pipelines can only be made public after authorization by an administrator, this is useful for organizations that desire to maintain strict control of broadly available pipelines.

Although integration with GitHub makes sharing and executing possible, pipelines can also be downloaded in NextFlow format for documentation, distribution and execution outside of DolphinNext. To allow users and administrators to make pipelines available across installations, DolphinNext supports pipeline import and export.

### Nested pipelines

Many pipelines share not just processes, but subcomponents involving several processes. For instance, BAM quality assurance is common to most sequence processing pipelines **(Figure 2. D)**. It relies on RseQC(40) and Picard(41) to create read quality control reports. To minimize redundancy, these modules can be encapsulated as pipelines and re-used as if they were processes. The pipeline designer module supports drag and drop of whole pipelines and in a similar way as it supports individual processes. Multiple pipelines such as RNA-Seq, ATAC-Seq, and ChIP-Seq can therefore have the same read quality assurance logic (**Figure S4**). Reusing complex logic by encapsulating it in a pipeline greatly simplifies and standardizes maintenance of data processing pipelines.

#### 3. Pipeline Executor

One of the most frustrating events in data processing is an unexpected pipeline failure. Failures can be the result of an error in the parameters supplied to the pipeline (e.g. an output directory with no write permissions, or incompatible alignment parameters) or because of computer system malfunctions. Restarting a process from the beginning when an error occurred at the very end of the pipeline can result in days of lost computing time.

DolphinNext provides a user interface to monitor pipeline execution in real time. If an error occurs the pipeline is stopped; the user, however, can restart the pipeline from the place where it stopped after changing the parameters that caused the error (**Figure 4**). Users can also assign pipeline runs to projects so that all pipelines associated with a project can be monitored together.

In addition to providing default values for options that are pipeline specific, administrators can provide default values for options common to all pipelines, such as resource allocation (e.g., memory, CPU, and time), and access level of pipeline results.

Specific features of pipeline running are:

##### Run status page

DolphinNext provides a “Run Status” page for monitoring the status of all running jobs that belong to the user or the groups to which the user belongs. This searchable table includes the name of the run, working directory, description, and status. Users can also access the run page of a specific run to make a change, terminate, resume, copy, delete, or re-execute.

##### Accessibility to all run logs

To monitor run specific details and troubleshoot failed runs, DolphinNext provides access to all application logs. Further, DolphinNext also gives access to NextFlow’s execution reports, which include the execution time, resource utilization (e.g. cpu or memory for each process), process duration time (both clock and CPU time) and memory utilization. In this way, users can optimize their computational resources by reducing unnecessary overhead based on the data available from past executions.

#### 4. Reports

Most processes in a pipeline produce interim data that is used by downstream processes. For example, in an RNA-Seq analysis pipeline, subtracting ribosomal reads prior to genomic alignment reduces processing time. This is done by aligning directly to the ribosomal RNA genes, and keeping reads with no matches for alignment to the genome or transcriptome(42). In addition to reducing compute time, the fraction of reads that align to ribosomal genes is an important metric to assess the technical efficiency of library preparation, specifically to measure how well ribosomal depletion worked. DolphinNext allows users to access and visualize interim pipeline results, such as intermediate alignments like alignment to the ribosomal RNAs, that are not part of the final result. Process designers can define any number of secondary outputs, which the DolphinNext report module can process and present in a user-friendly format. While there is an integrated table viewer for tabular data, these data can also be seamlessly loaded and analyzed using an embedded DEBrowser, an interactive visualization tool for count data(30). PDF and HTML files can be visualized within DolphinNext or downloaded for offline analysis (**Figure 5**).

**Figure 5.**
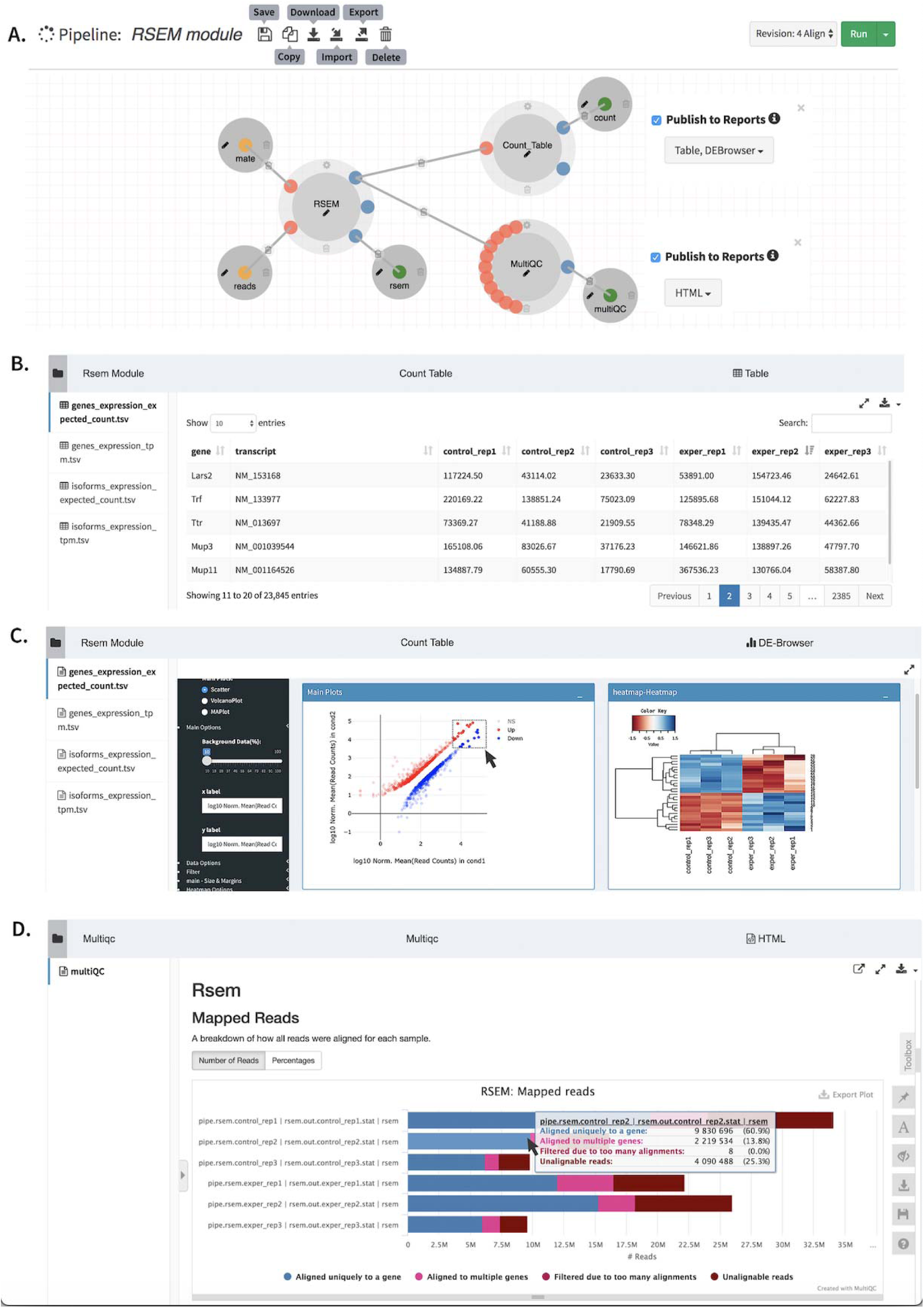
**A.** RSEM module which involves Count_Table to summarize sample counts into a consolidated count table. This process reports the results with a table or upload the count table to embedded DEBrowser(30), **B.** Count table report **C.** MultiQC(43) report to summarize numerous bioinformatics tool results, and **D.** Embedded DEBrowser(30) module for interactive differential expression analysis.

For more flexible reports, DolphinNext supports an R-markdown defined output. Process designers can include a custom R markdown script specifically designed to handle and visualize the process output. Pipeline designers can make this report available to users for interactive analysis of the process results (Figure 6).

**Figure 6.**
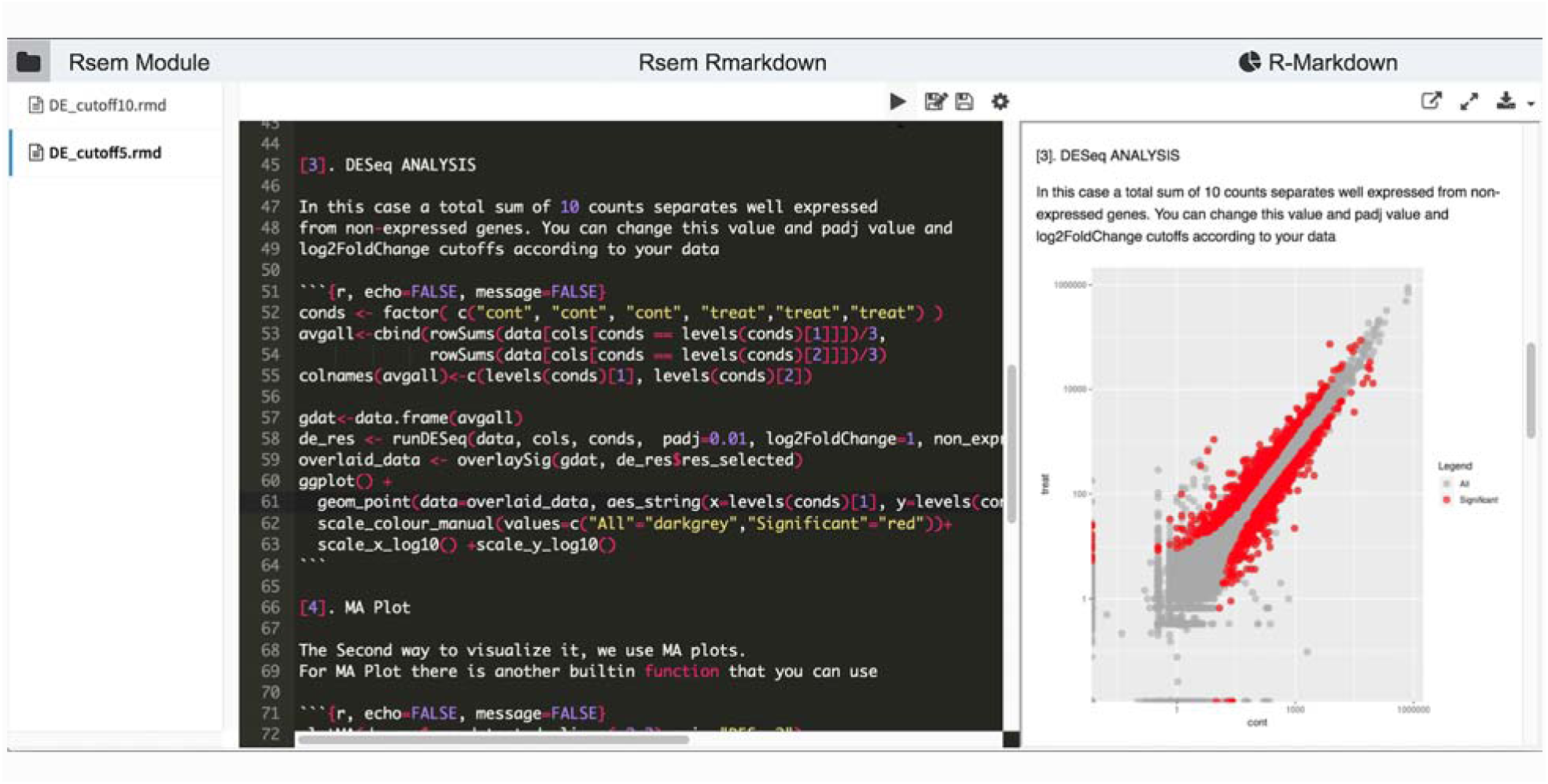
R markdown report for RNA-Seq analysis. Users can adapt this template according to their needs.

## RESULTS

We highlight the flexibility of DolphinNext’s pipeline builder by providing a detailed description of end-to-end RNA-Seq and ChIP-Seq/ATAC-Seq processing pipelines. Each of these pipelines provides unique features not found in other publicly available workflows such as: 1. Extensive support for pre-processing (read trimming by quality or adapter sequence, iterative removal of specific RNA or DNA species, such as rRNA and repetitive sequences), 2. Support for different aligners and quantification methods in the processing steps, and 3. An extensive reports and quality control checks in the post-processing steps.

### 1. RNA-Seq Pipeline (Figure S1)

All sequence processing pipelines take one or several fastq input files. This pipeline, like all other high-throughput sequencing processing pipelines (see below for other examples), first performs **data preparation**, a step that consists of basic read quality control and filtering actions **(Figure S2)**: read quality reports, read quality filtering, read quality trimming, adapter removal. After these quality control steps, the RNA-Seq pipeline offers the user the option to align, filter out, and/or estimate the abundance of both standard and predefined sets of genomic loci (e.g. rRNAs, miRNAs, tRNAs, piRNAs, snoRNAs, ERCC(44), mobile elements). Before any such alignment, reads may be trimmed to a desired length for optimal alignment, especially if quality issues in the 3’ or 5’ ends of the reads are suspected (e.g. to align miRNA, or tRNA sequences). Once the data preparation phase of the pipeline is complete, the pipeline produces quality reports including FastQC reports(45) and information about the fraction of reads aligned to each of the genomic loci selected by the user. In its current implementation, data preparation relies on FastQC(46), adapter removal using cutadapt(47), 5’ and 3’ trimming, quality removal using trimmomatic(48), and Bowtie2(49) for alignments against selected regions or transcripts (**Figure S3**).

Only reads that pass all filters in the data preparation stage are kept for later steps. To estimate expression levels, the RNA-Seq pipeline uses RSEM(39) which aligns reads to a predefined set of transcripts. The user can use available transcript sets (i.e Ensemble(50), GENCODE(51, 52), RefSeq(53)) or upload their own. The RNA-Seq pipeline also allows the user to align reads against the genome using splicing aware alignment algorithms and generate a genome browser viewable file to visually assess genomic contamination and library quality. To do so, the user may choose between any or all of the most commonly used aligners: STAR(54), Hisat2(55) and Tophat2(56). Resulting alignments are then processed to generate genome browser friendly formats: bigwig (for UCSC genome browser(32) or TDF (for the integrative genome viewer (IGV, (31)).

If the user opted to perform genomic alignments, the pipeline reports overall quality metrics such as coverage and the number of mapped reads to different genomic and transcriptomic regions (**Figure S4**). These reports rely on Picard’s CollectRNASeqMetrics program(41) and the RSeQC(40) program.

Finally, the RNA-Seq pipeline returns a quantification matrix that includes the estimated counts and transcripts per million (TPM) based on RSEM(39) or by simply counting reads using featureCounts(57) for each gene and/or for each annotated isoform. These matrices are used as inputs for differential gene expression analysis and can be uploaded directly to an embedded instance of our DEBrowser(30) software, which allows interactive exploration of the resulting data(30).

### 2. ATAC-Seq and ChIP-Seq pipeline (Figure S4-S5)

DolphinNext offers pipelines to process libraries for the analysis of ChIP-Seq and ATAC-Seq data. These two pipelines share most processes and only differ at a few very specific points. They also share all initial data preparation steps with the RNA-Seq pipeline, such that both rely on the very same processes for read filtering, read quality reporting and alignment to desired genomic locations to quantify and filter out the reads mapped to repeat regions or any other loci of interest. Filtering out reads mapping to a predefined set of loci can dramatically speed up the genome-wide alignment that follows.

After data processing, reads are aligned to the genome using a short read aligner such as Bowtie2(49). For large data files, such as those typically obtained from ATAC-Seq, alignments can be sped up by splitting the files into smaller chunks that are aligned in parallel. The choice to split and the chunk sizes should be determined by the user based on the specific computing environment. By default, the pipeline creates smaller files that each have 5 million reads. After alignments of each read chunk, the results are automatically merged with the samtools(58) merge function. The pipeline then allows users to estimate and remove PCR duplicates using the Picard’s mark duplicates(41) function.

For ATAC-Seq, the pipeline calls accessible chromatin regions by estimating the Tn5 transposase cut sites through a read extension process that has been shown to more accurately reflect the exact position that was accessible to transposase(42, 59). Once each read has been shortened, peaks are called identically in both the ChIP-Seq and ATAC-Seq using MACS2(60).

When processing several samples together, the ATAC and ChIP pipelines provide consensus peak calls by merging all peaks individually called in each sample using Bedtools(61). The number of reads in each peak location are then quantified using Bedtools(61) coverage function. As a result, ATAC-Seq and ChIP-Seq pipelines also generate a matrix that has the count values for each peak region and samples. This matrix can be uploaded directly to the embedded version of DEBrowser(30) to perform differential analysis or downloaded to perform other analysis. Finally, to determine motifs under the peaks in Chip-Seq pipeline, there is also a motif discovery support using HOMER (http://homer.salk.edu/homer/index.html).

## DISCUSSION

DolphinNext is a powerful workflow design, execution, and monitoring platform. It builds on top of recent technological advances such as software containerization and the Nextflow engine to address current data processing needs in high-throughput biological data analysis. Its ability to run anywhere, leverage the computing infrastructure of the institution, and provide an intuitive user interface makes it suitable for both small, large, and complex projects.

Reproducing third party analyses that involve many different programs, each with custom parameters, is a tremendous challenge. We have put special emphasis on the reproducibility of pipelines. To enable this, DolphinNext investigators can distribute their processing pipelines as containers that can run as stand-alone applications including the proper versions of all software dependencies. Further, DolphinNext supports pipeline versioning where users can create and tag a pipeline version with a unique identifier (uid) that includes the general pipeline “graphical representation”, the executable nextflow script, as well as the exact software versions and parameters used when run on a specific dataset.

Users with different computational skills can use DolphinNext to increase productivity. Casual users can rely on previously defined and tested pipelines to process their datasets. Investigators can easily distribute their processing pipeline along with the data for others to reproduce their analyses. Investigators with access to parallel computing systems but without strong computational background can use DolphinNext to optimally utilize their computing environment. For example, DolphinNext pipelines can automatically split large sequence data files into smaller “chunks” that can be aligned in parallel across hundreds of cluster nodes if such infrastructure is available. Finally, bioinformaticians can easily create new processes and integrate them into custom robust and reproducible pipelines.

DolphinNext offers a highly modular platform. Though this manuscript only focuses on the power of DolphinNext for data processing, we have also integrated two different downstream analysis tools (DEBrower(30) and Rmarkdown) that take the count matrices output from the data processing steps directly into analysis and data exploration steps. Furthermore, DolphinNext’s modular architecture allows for easy integration of any custom data analysis and visualization tool. As such DolphinNext is meant to provide a basic platform to standardize processing and analysis across institutions.

## AVAILABILITY

DolphinNext is an open source collaborative initiative available in the GitHub repository (https://github.com/UMMS-Biocore/dolphinnext). A documentation, quick start guides and video tutorials are available in readthedocs. (https://dolphinnext.readthedocs.org). A docker image to run DolphinNext is in the GitHub repository (https://github.com/UMMS-Biocore/dolphinnext-studio). The web server is available in https://dolphinnext.umassmed.edu. Example pipelines are publicly available in the GitHub repository at https://github.com/dolphinnext.

## ACKNOWLEDGMENTS

We would like to thank Phil Zamore and members of his lab for their advice and suggestions and Athma Pai, Alan Derr, Rachel Murphy, Kyle Gellatly, and Laney Zuerlein for constructive suggestions, to all members of the Garber Lab for testing and improving pipelines and for their comments on the manuscript. We would also like to thank Craig C. Mello and his lab to initiate the project.

## AUTHORS’ CONTRIBUTIONS

OY and AK implemented project. AK and MG supervised the project and wrote the manuscript. OT and ARO helped developing pipeline builder visualization module. All authors have read and approved the final manuscript.

