## Supplemental Figures for "DolphinNext: A graphical user interface for creating, deploying and executing Nextflow pipelines"

### SUPPLEMENTARY FIGURES

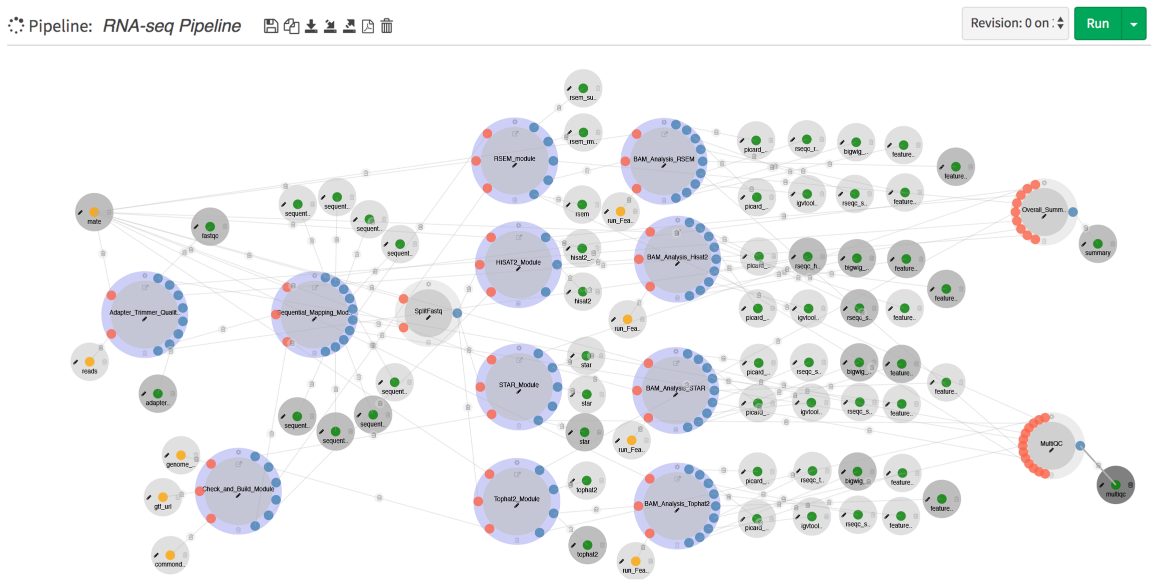


**Figure S1.** RNA-Seq Pipeline

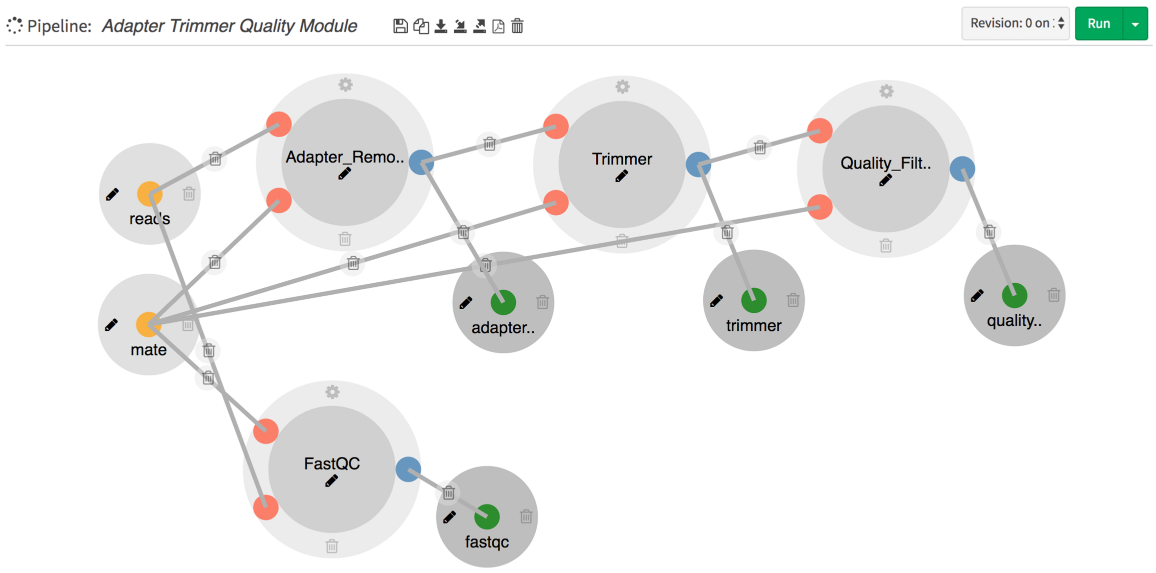


**Figure S2.** Adapter Removal, Trimmer and Quality Filtering Module

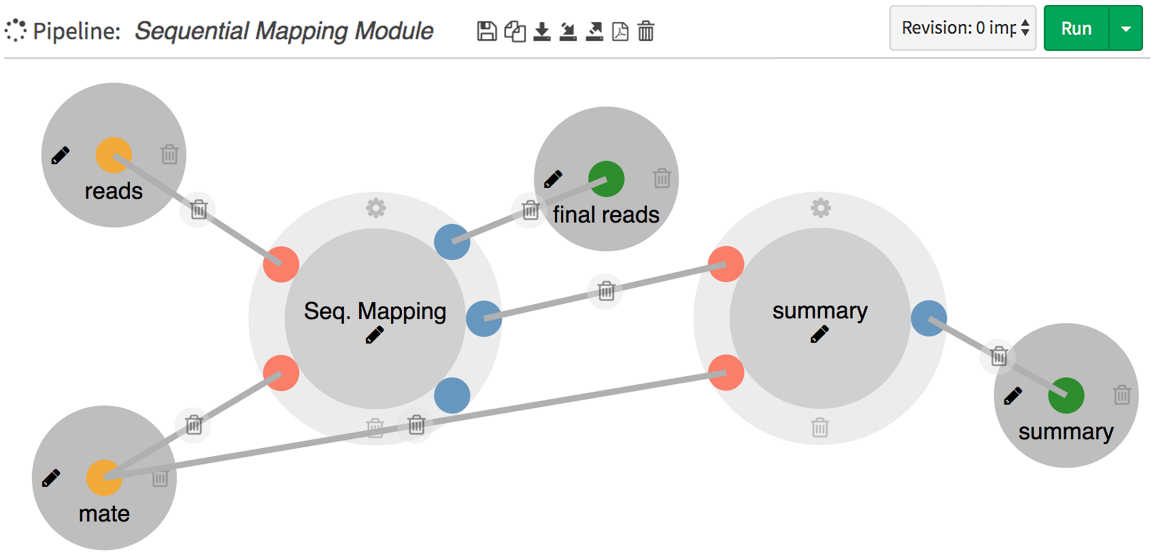


**Figure S3.** Sequential Mapping Module

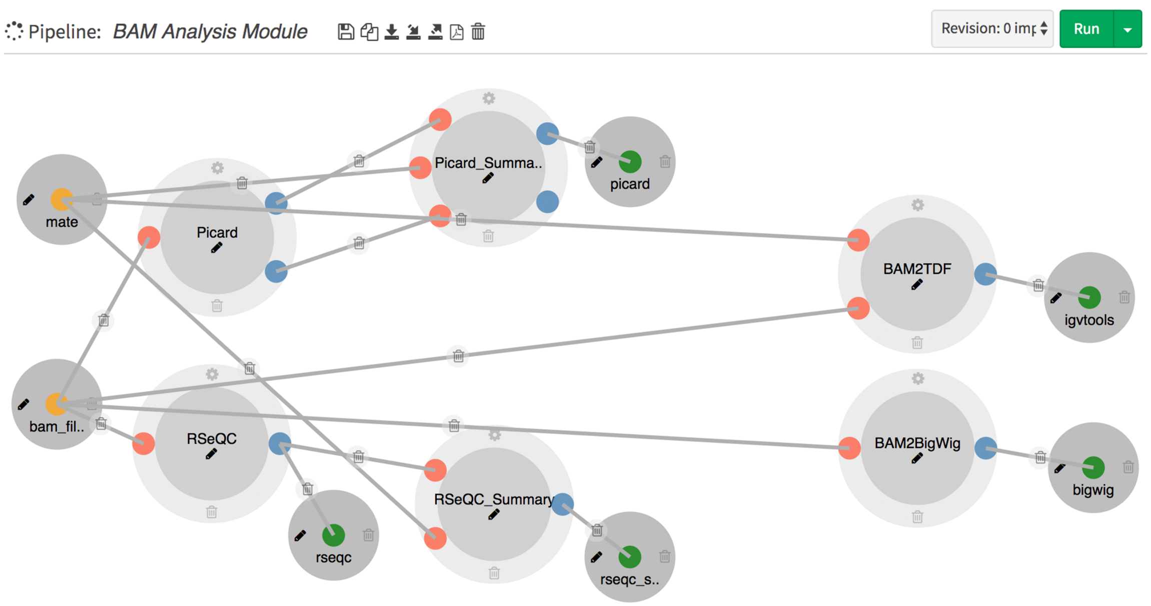


**Figure S4.** BAM Analysis Module

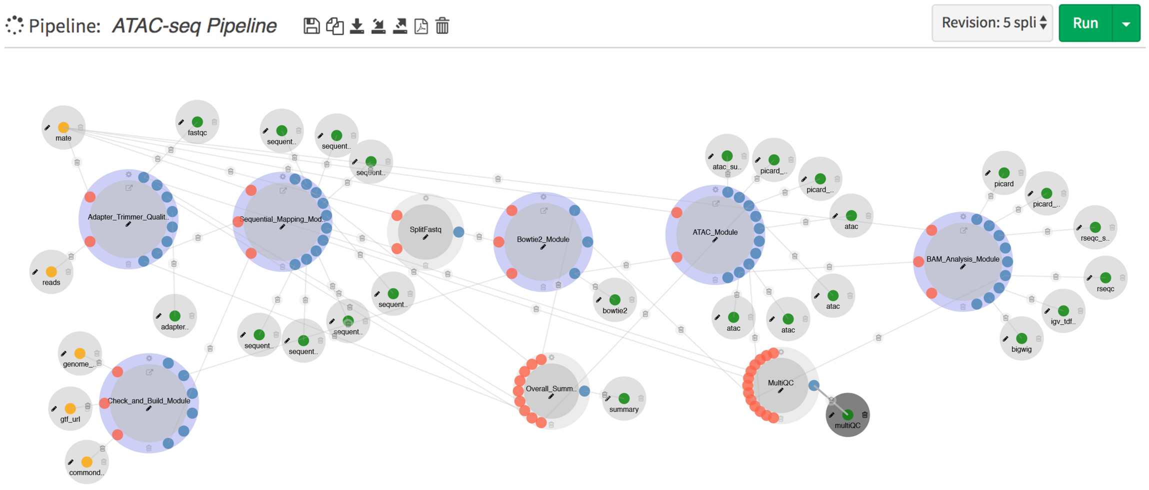


**Figure S5.** ATAC-Seq pipeline

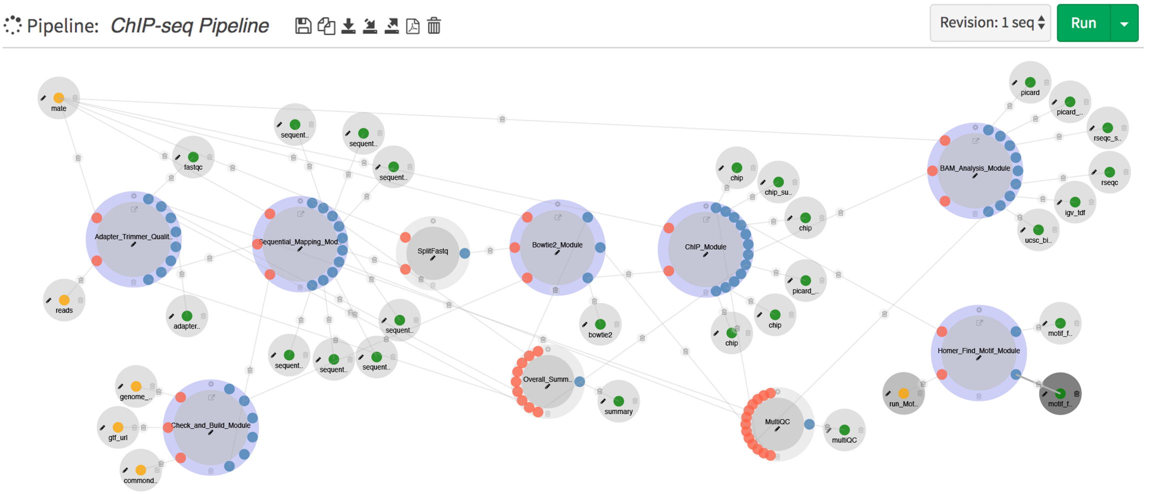


**Figure S6.** ChIP-Seq pipeline
